# Cortical and thalamic connectivity of posterior parietal visual cortical areas PPc and PPr of the domestic ferret (*Mustela putorius furo*)

**DOI:** 10.1101/491993

**Authors:** Leigh-Anne Dell, Giorgio M. Innocenti, Claus C. Hilgetag, Paul R. Manger

## Abstract

The present study describes the ipsilateral and contralateral cortico-cortical and cortico-thalamic connectivity of the parietal visual areas PPc and PPr in the ferret using standard anatomical tract-tracing methods. The two divisions of posterior parietal cortex of the ferret are strongly interconnected, however area PPc shows stronger connectivity with the occipital and suprasylvian visual cortex, while area PPr shows stronger connectivity with the somatomotor cortex, reflecting the functional specificity of these two areas. This pattern of connectivity is mirrored in the contralateral callosal connections. In addition, PPc and PPr are connected with the visual and somatomotor nuclei of the dorsal thalamus. Numerous connectional similarities exist between the posterior parietal cortex of the ferret (PPc and PPr) and the cat (area 7 and 5), indicative of the homology of these areas within the Carnivora. These findings highlight the existence of a fronto-parietal network as a shared feature of the organization of parietal cortex across Euarchontoglires and Laurasiatherians, with the degree of expression varying in relation to the expansion and areal complexity of the posterior parietal cortex. This observation indicates that the ferret is a potentially valuable experimental model animal for understanding the evolution and function of the posterior parietal cortex and the fronto-parietal network across mammals. The data generated will also contribute to the Ferretome (www.ferretome.org) connectomics databank, to further cross-species analyses of connectomes and illuminate wiring principles of cortical connectivity across mammals.

## Introduction

The posterior parietal cortex is a multisensory cortical territory, of varying size within and between different mammalian lineages, situated between the unimodal somatosensory and visual cortical territories (e.g. Hyvärinen, 1982; Caminiti, Innocenti, & Battaglia-Mayer, 2015; Kaas & Stepniewska, 2016; Goldring & Krubitzer, 2017; Kaas, Qi, & Stepniewska, 2017). This region of cortex is involved in many functions including sensory perception, motor activity and behavioural modulation (see Hyvärinen, 1982 for an extensive review). Studies of the parietal region of the cortex have mostly been undertaken in primates, due to the expansion of this region in the primate lineage and especially so in humans (Hill, Inder, Neil, Diesker, Harwell, & Van Essen, 2010), the importance of this region in visually-guided motor tasks (Creem-Regehr, 2009), and the small size of this cortical region in other members of the Euarchontoglires (Kolb & Walkey, 1987; Wallace, Ramachandran, & Stein, 2004; Remple, Reed, Stepniewska, & Kaas, 2006; Remple, Reed, Stepniewska, Lyon, & Kaas, 2007; Goldring & Krubitzer, 2017). In primates, and especially in humans, this cortical territory is thought to be parcellated into many different functional regions or areas (e.g. Seltzer & Pandya, 1980; Lewis & Van Essen, 2000a,b; Nelson et al., 2010; Mars, Sallet, Schüffelgen, Jbabdi, Toni, & Rushworth, 2011; Stepniewska, Friedman, Gharbawie, Cerkevich, Roe, & Kaas, 2011; Kaas, Qi, & Stepniewska, 2017), and it is heavily interconnected with motor, premotor and other cortical regions (e.g. Felleman & Van Essen, 1991; Bullmore & Sporns, 2005; Cloutman & Lambon Ralph, 2012; Kaas & Stepniewska, 2016; Kaas et al., 2017). In the smallest-brained primate where posterior parietal cortex has been extensively studied (Galago garnetti), this cortical territory has been subdivided into three distinct areas − rostral (visuomotor), caudal (visual) and medial (possibly visuomotor) (Fang, Stepniewska, & Kaas, 2005; Stepniewska, Fang, & Kaas, 2009a; Stepniewska, Cerkevich, Fang, & Kaas, 2009b; Kaas & Stepniewska, 2016), with the rostral area being heavily interconnected with the motor and pre-motor cortex (Stepniewska et al., 2011; Kaas & Stepniewska, 2016; Kaas et al., 2017).

In contrast to the studies undertaken in mammals belonging to the Euarchontoglires, studies of species within the other major Eutherian mammal radiation, the Laurasiatheria, are more limited, and only found in species of the order Carnivora. Brodmann (1909) provided an architectonic study of the posterior parietal cortex in the kinkajou, but the majority of studies have focused on the cat (Heath & Jones, 1971; Marcotte & Updyke, 1982; Symonds & Rosenquist, 1984; Olson & Lawler, 1987; Pigarev & Rodionova, 1998). In addition, two studies have examined the posterior parietal cortex of the ferret (Manger, Masiello, & Innocenti, 2002b; Homman-Ludiye, Manger, & Bourne, 2010). These studies, through architecture, physiological mapping and limited connectional studies, have revealed the presence of two distinct cortical areas within the ferret posterior parietal cortex, a caudal unimodal visual area and a rostral bimodal somatovisual area.

The ferret posterior parietal cortex appears to be organized in a manner quite similar to that observed in the galago, although the mosaic or fractured maps of sensory surfaces found in the galago and other primates (Kaas & Stepniewska, 2016; Kaas et al., 2017) was not observed in the ferret (Manger et al., 2002b). Despite this seemingly similar organization, it must be noted that the last common ancestor of the ferret and galago occurred 85-95 million years ago (Kaschube, Schnabel, Löwel, Coppola, White, & Wolf, 2010; Kiel et al., 2012; Kaas et al., 2017). Given that the non-primate Euarchontoglires only have a small posterior parietal cortical region (Kaas & Stepniewska, 2016; Goldring & Krubitzer, 2017; Kaas et al., 2017), it is highly unlikely that the cortical areas in the ferret and galago can be considered homologous − the relative expansion of the posterior parietal cortex in the different lineages (primate and carnivore) being potentially independent occurrences. This observation then raises the question of whether these areas, if not directly homologous, are analogous. The extensive fronto-parietal network associated with visually guided behaviour is a clear trait of the posterior parietal cortex in the primates (Kaas & Stepniewska, 2016; Kaas et al., 2017). If such a fronto-parietal network were to exist in the ferret, the case for analogy of these regions in the different species would be quite strong, and indeed be informative about the baseline organization of the posterior parietal cortex in early Eutherian mammals, and the potential evolutionary pathways and constraints that have led to the organization of the posterior parietal cortex in the different major Eutherian mammal lineages (Euarchontoglires and Laurasiatheria). To this end, the current study examined the corticocortical and thalamic connectivity of the posterior parietal cortical areas **PPc** and **PPr** of ferret in order to determine whether an extensive fronto-parietal network is present.

## Material and Methods

### Surgical procedure and tracer injections

Eight adult female ferrets (Mustela putorius furo), weighing between 600g and 1000g, were used in this current study (four injection sites per cortical area). The experiments were conducted according to the Swedish and European Community guidelines for the care and use of animals in scientific experiments. All animals were initially anesthetized with i.m. doses of ketamine hydrochloride (Ketalar, 10mg/kg) and medetomidin hydrochloride (Domitor, 0.08mg/kg), supplemented with atropine sulphate (0.15mg/kg) and placed in a stereotaxic frame. A mixture of 1% isoflurane in a 1:1 nitrous oxide and oxygen mixture was delivered through a mask while the animal maintained its own respiration. Anesthetic level was monitored using the eye blink and withdrawal reflexes, in combination with heart rate measurement. The parietal cortex was exposed under aseptic conditions and in each animal numerous (but fewer than 20) electrophysiological recordings were taken to ensure correct placement of the tracer within a specific cortical area (Manger et al., 2002b). Approximately 500 nl of tracer (biotinylated dextran amine, BDA 10 k, 5% in 0.1 M phosphate buffer; Molecular Probes) was delivered at each injection site using a Hamilton microsyringe (Figs. 1, 2a, 2b).

**Figure 1:**
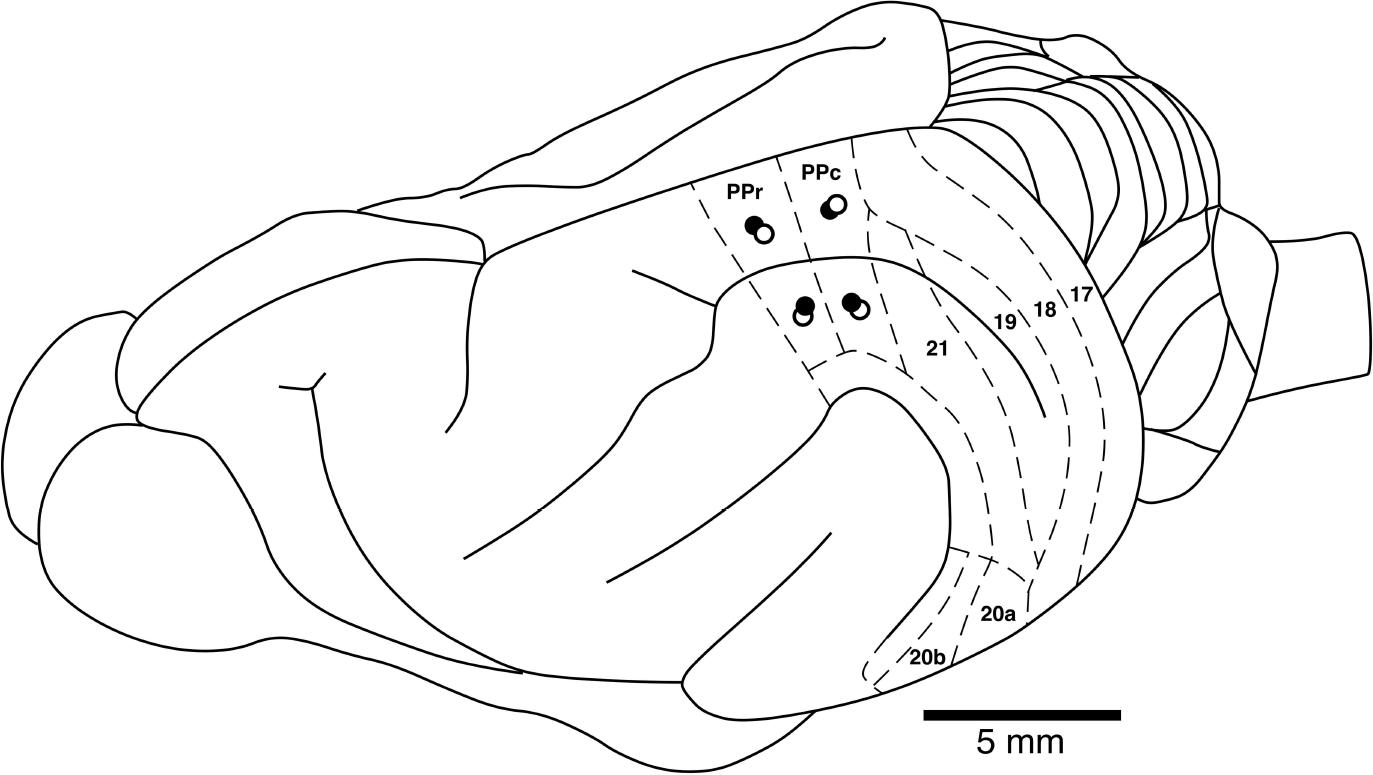
Locations of the injection sites in the parietal cortical areas analyzed in the current study in relation to many of the known boundaries of visual cortical areas in a dorsolateral view of the ferret brain. Closed circles represent the injections sites where the brain was sectioned in a coronal plane. Open circles represent the injections sites where the cerebral cortex was manually semi-flattened for analysis. See list for abbreviations.

**Figure 2:**
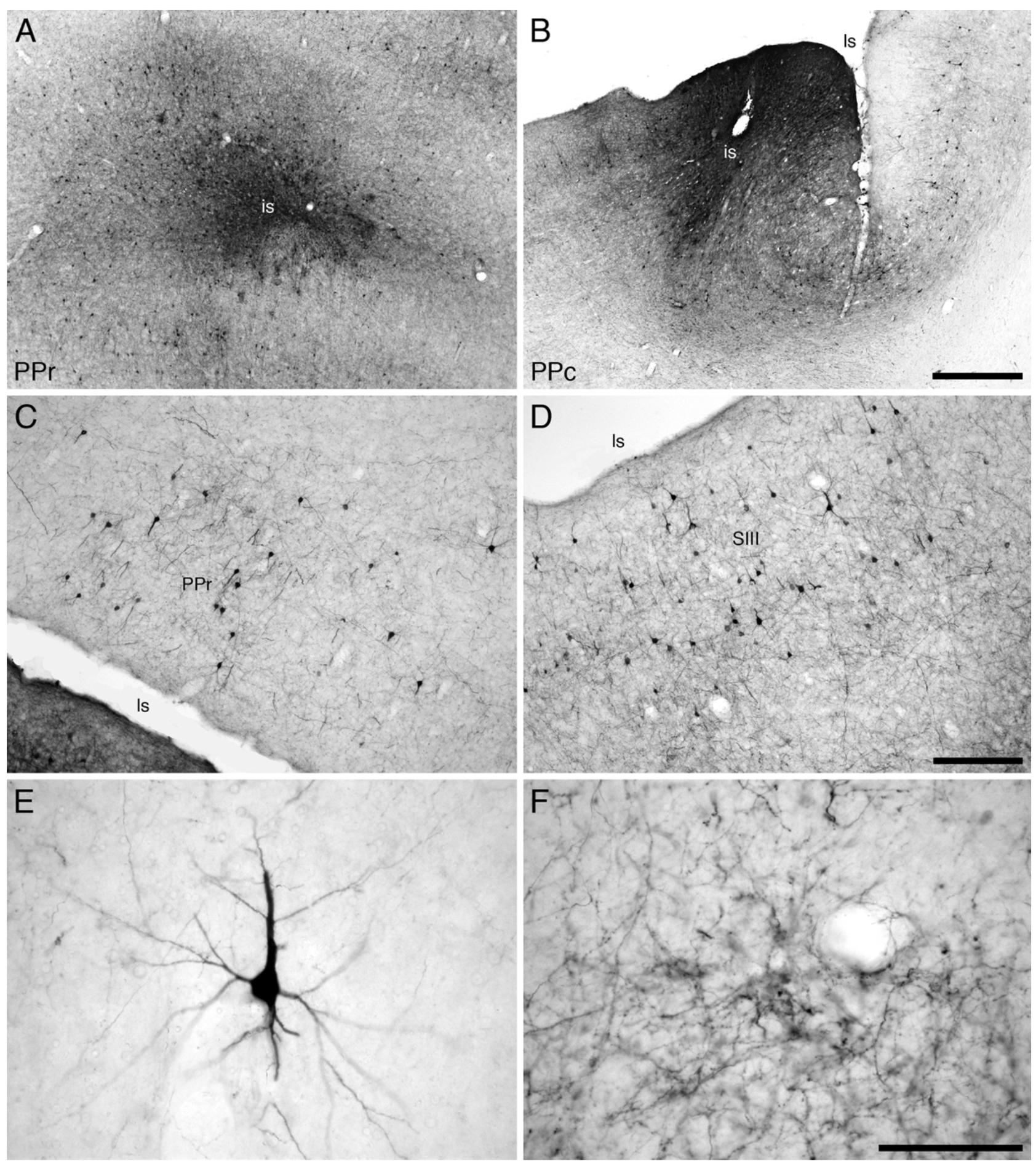
Photomicrographs showing examples of injection sites (**a**, **b**) and labeled cells and axons terminals (c-f) that were analyzed in the current study. (**a**) Injection site (**is**) in area **PPr** of a semi-flattened cerebral cortex showing the spread of tracer around the injection site, indicating the areal specificity of the injections made in the current study. (**b**) Injection site (**is**) in area **PPc**from a coronal section through the cerebral cortex, showing the spread of tracer and the limitation of the injection site to the cerebral cortex. Labeled cells and terminals in the medial portion of area **PPr** (**c**, medial to the lateral sulcus, **Is**) and **SII** I (**d**, lateral to the lateral sulcus) following transport from an injection site in area **PPr**. (**e**) High magnification image of a retrogradely labeled cell in area **SII**following transport from the injection site depicted in **a**. (**f**) High magnification image of anterogradely labeled axons in area **21** following transport from the injection site depicted in **a**. Scale bar in **b** = 500 μm and applies to **a** and **b**. Scale bar in **d** = 250 μm and applies to **c** and **d**. Scale bar in **f** = 100 μm and applies to **e** and **f**. In images **a**, **c**, **d**, and **f**, the midline of the brain is to the top of the image and rostral to the left. In image **b**, medial is to the right and dorsal to the top. In image **e** the pial surface has been rotated to the top of the image.

After the completion of the injections, a soft contact lens was cut to fit over the exposed cortex, while the retracted dura mater was pulled over the contact lens and the excised portion of bone repositioned and held in place with dental acrylic. The temporal muscle was reattached using surgical glue and the midline incision of the skin sutured. Antibiotics were administered (Terramycin, 40 mg/kg, daily for 5 days) and these animals were given a 2-week recovery period to allow for tracer transport. At the end of this period, the animals were euthanized with a lethal dose of sodium pentobarbital (80 mg/kg, i.p.) and perfused intracardially, initially with a rinse of 0.9% saline (4°C, 500ml/kg), followed by fixation with 4% paraformaldehyde in 0.1M phosphate buffer (4°C, 1000 ml/kg).

### Sectioning and staining procedures

The brains were removed from the skull and post-fixed overnight in 4% paraformaldehyde in 0.1M phosphate buffer and then transferred to a 30% sucrose solution in 0.1M phosphate buffer (4°C) and allowed to equilibrate. The brains were either: (1) frozen in dry ice and sectioned at 50 μm on a freezing microtome in a coronal plane (4 cases, two for each of the parietal cortical areas, one in the medial and one in the lateral portion of each area) for a one in four series for Nissl (cresyl violet), myelin (Gallyas, 1979), cytochrome oxidase (Carroll & Wong-Riley, 1984) and BDA; or (2) cryoprotected and the cerebral cortex dissected away from the remainder of the brain and the dorsolateral surface semi-flattened (4 cases, two for each of the parietal cortical areas, one in the medial and one in the lateral portion of each area) between two glass slides, frozen onto the cold microtome stage and sectioned parallel to the semi-flattened surface at 50 μm for a one in two series for BDA and cytochrome oxidase. For BDA tracer visualization, the sections were incubated in 0.5% bovine serum albumin in 0.05M Tris buffer for 1h, followed by incubation in an avidin-HRP solution for 3h. A 10 min pre-incubation in 0.2% NiNH_4_SO_4_ preceded the addition of H_2_O_2_ (200 μl/l) to this solution, at which time the sections were monitored visually for the reaction product. To stop the reaction, the sections were placed in 0.05 M Tris buffer. All sections were mounted on 0.5% gelatine coated slides, dehydrated in a graded series of alcohols, cleared in xylene and coverslipped with Depex mounting medium. All injection sites resulted in robust anterograde and retrograde transport of tracer in both the cerebral cortex (Fig. 2c-f) and the visual and motor portions of the dorsal thalamus (Fig. 3).

**Figure 3:**
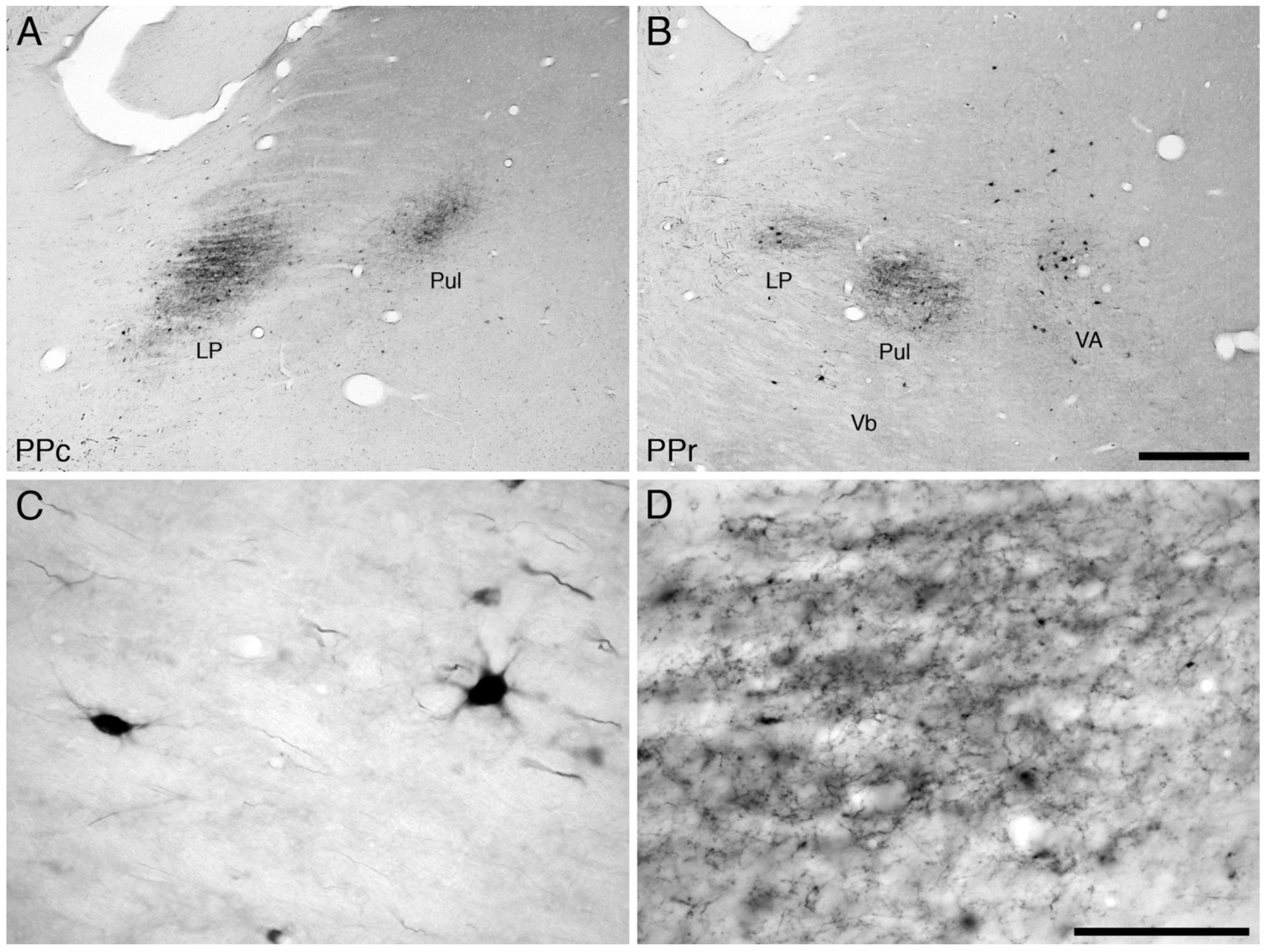
Photomicrographs showing examples of retrogradely labelled cells and anterogradely labelled axons in the visual and motor thalamus of the ferret following injections into the parietal visual areas **PPc**(**a**), and **PPr**(**b**). (**c**) High magnification image of retrogradely labelled cells in the ventrobasal complex (**Vb**) following transport from the injection site in area **PPr**. (**d**) High magnification image of anterogradely labelled axons in the pulvinar nucleus following transport from the injection site in area **PPc**. Scale bar in **b**= 500 μm and applies to **a** and **b**. Scale bar in **d**= 100 μm and applies to **c** and **d**. In all images dorsal is to the top and medial to the right. See list for abbreviations.

### Qualitative and quantitative analysis

For qualitative analysis, the stained sections were examined under low and high power magnification using a light microscope to determine in which sections through the cortex labelled cell bodies and terminals were present. Under low power stereomicroscopy using the flattened sections, the edges of each section were drawn with the aid of a camera lucida, and the location of the injection site marked. Areal borders were delineated and drawn using the cytochrome oxidase stained sections. The sections reacted for BDA were then matched to these drawings and the locations of the individual retrogradely labelled cells plotted and regions of anterogradely labelled axonal terminals demarcated. The drawings were scanned and redrawn using a Canvas X Pro 16 drawing program (ACD Systems International Inc., USA). Digital photomicrographs were captured using a Zeiss Axioskop and the Axiovision software. No pixilation adjustments or manipulation of the captured images were undertaken, except for contrast and brightness adjustment using Adobe Photoshop 7.

To quantify the retrograde BDA labelling per injection site, and control for variance in the size of the injection, a fraction of labelled neurons (N%, Table 1) was calculated using the formula:

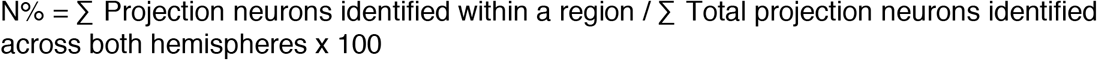

Where cell bodies could be clearly identified, neurons within the injection halo were counted. Furthermore, cell bodies located on boundary lines between areas were only accounted for in the region where the most overlap was present.

**Table 1:**
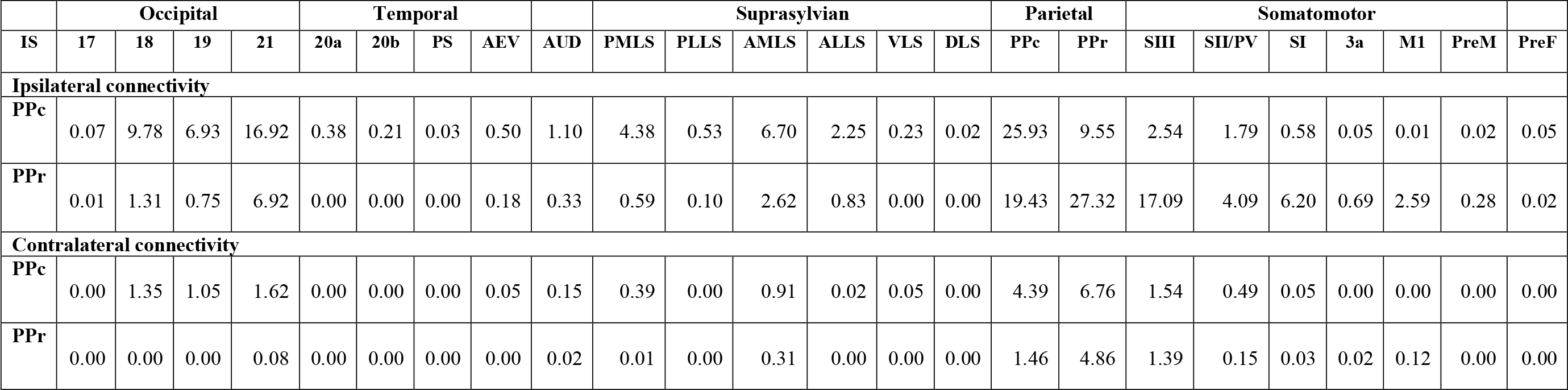
Average fraction of retrogradely labelled neurons (N%), in the various visual cortical areas of the ipsilateral and contralateral hemispheres of the ferret following four separate injections of biotinylated dextran amine (BDA) into the parietal visual areas **PPc** and **PPr**. See list for abbreviations. **IS** - injection site

## Abbreviations

**3a,** rostral, or proprioceptive, somatosensory area
**17,** primary visual cortex
**18,** second visual cortical area
**19,** third visual cortical area
**20a,** temporal visual area a
**20b,** temporal visual area b
**21,** fourth visual cortical area
**A,** A lamina of the lateral geniculate nucleus
**A1,** A1 lamina of the lateral geniculate nucleus
**AI,** primary auditory cortex
**AAF,** anterior auditory field
**ADF,** anterior dorsal auditory field
**AEV,** anterior ectosylvian visual area
**AMLS,** anteromedial lateral suprasylvian visual area
**ALLS,** anterolateral lateral suprasylvian visual area
**AUD,** auditory cortex
**AV,** anterior ventral nucleus of the dorsal thalamus
**AVF,** anterior ventral auditory field
**C,** C lamina of the lateral geniculate nucleus
**DLS,** dorsal lateral suprasylvian visual cortical area
**is,** injection site
**LP,** lateral posterior nucleus of the dorsal thalamus
**ls,** lateral sulcus
**M1,** primary motor cortex
**MIN,** medial intralaminar nucleus of the lateral geniculate nucleus
**OB,** olfactory bulb
**OT,** olfactory tract
**P,** perigeniculate lamina of the lateral geniculate nucleus
**PIR,** piriform cortex
**PMLS,** posteromedial lateral suprasylvian visual area
**PLLS,** posterolateral lateral suprasylvian visual area
**Po,** posterior nucleus of the dorsal thalamus
**PPc,** posterior parietal caudal cortical area
**PPr,** posterior parietal rostral cortical area
**PreF,** prefrontal cortical region
**PreM,** premotor cortical region
**PS,** posterior suprasylvian visual cortical area
**PSF?,** potential posterior suprasylvian auditory field
**Pul,** pulvinar nucleus of the dorsal thalamus
**SI,** primary somatosensory area
**SII/PV,** second somatosensory area/parietoventral somatosensory area
**SIII,** third somatosensory area
**SMA?,** potential supplementary motor area
**VA,** ventral anterior nucleus of the dorsal thalamus
**Vb,** ventrobasal complex of the dorsal thalamus
**VLS,** ventral lateral suprasylvian visual area
**VL,** ventral lateral nucleus of the dorsal thalamus
**VP?,** ventral posterior ectosylvian region.

## Results

In the current study, anatomical tract tracer injections were placed into the medial and lateral aspects of the caudal and rostral posterior parietal cortical areas (**PPr** and **PPc**) of the ferret, in order to examine the cortico-cortical and cortico-thalamic connections of these two parietal cortical areas. Despite minor quantitative differences being observed between the injections placed in the lateral and medial aspects of both **PPc** and **PPr**, the overall distribution of labelled connections was identical between the lateral and medial portions of each cortical area. Thus, the description of connectivity provided herein has grouped all four injections in each area for brevity.

It was evident in all cases that there was more connectivity in the hemisphere ipsilateral to the injection site than in the contralateral hemisphere, and that this label was clustered in its distribution (Table 1, Fig. 4). In all cases, ipsilateral connectivity was strongest within the parietal visual areas, the somatosensory areas, and the suprasylvian visual area **AMLS**. Moderate connectivity strength was observed the occipital visual areas and the remaining suprasylvian visual areas, while weak connectivity was observed in the temporal visual areas and frontal cortex. Contralateral connectivity was strongest in the regions homotopic to the injection site, while connectivity with the dorsal thalamus was found primarily in specific visual and motor nuclei.

**Figure 4:**
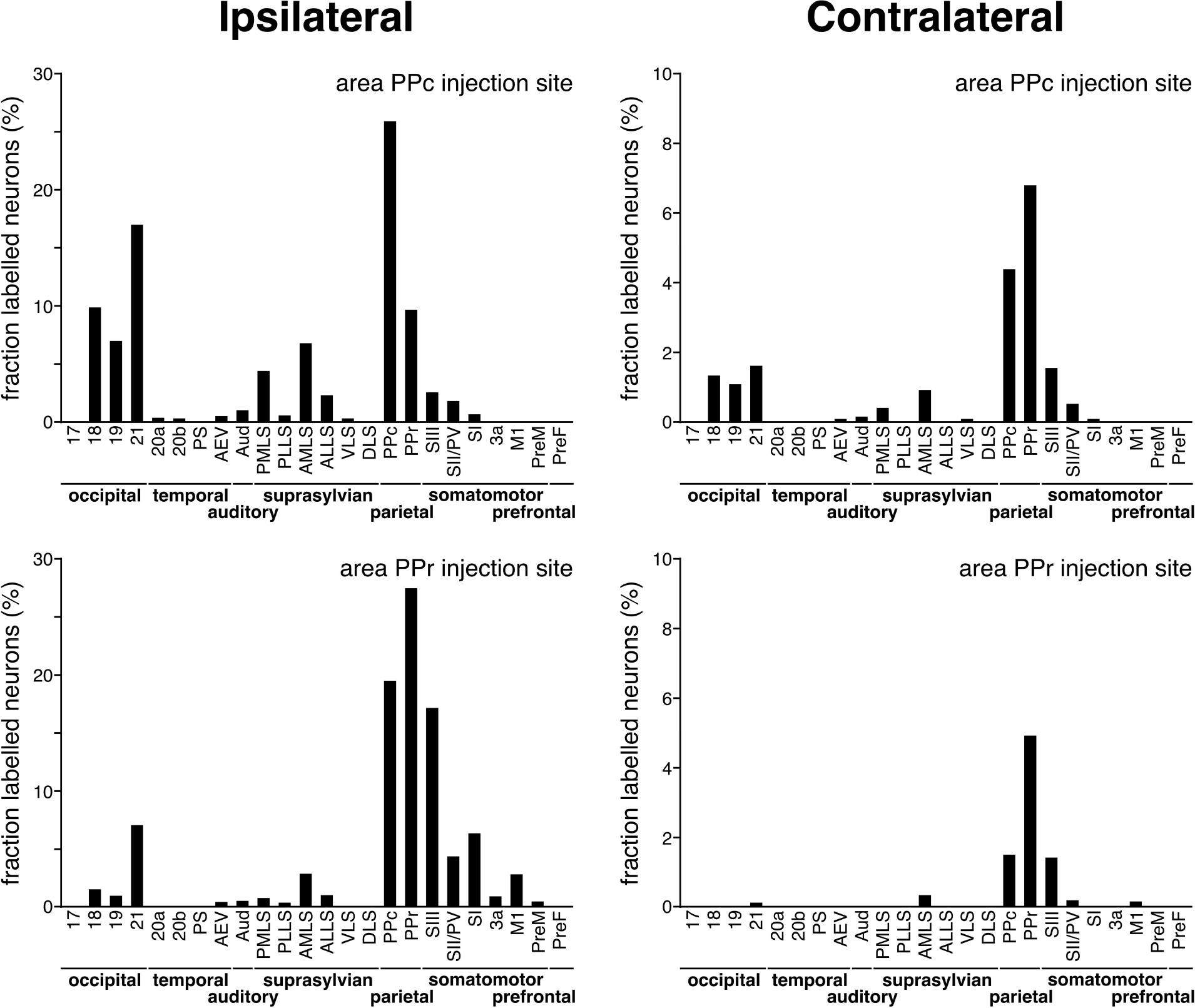
Graphs depicting the quantitative assessment of retrograde connectivity strength within and between cortical areas in the cerebral hemispheres ipsilateral (**left column**) and contralateral (**right column**) to the injection sites made in areas **PPc** (**top two graphs**) and **PPr**(**bottom two graphs**). The values are expressed in percentages, being the fraction of labelled neurons occurring in each cortical area. See list for abbreviations. Note that the majority of retrogradely labelled cells are found within the cortical area injected, and that both parietal areas are connected to each other, both ipsilaterally and contralaterally.

### Connectivity of PPc

Injection of tracer into **PPc** revealed widespread connectivity throughout the vast majority of the ipsilateral cerebral cortex (Fig. 5), with connections observed in parietal, occipital, suprasylvian and temporal visual regions, throughout the somatomotor and prefrontal regions, the auditory and multimodal cortex of the anterior and middle ectosylvian gyrus, as well as with the cortex overlying the claustrum. The only region of the cerebral cortex devoid of any connectivity was the posterior ectosylvian gyrus (Fig. 5). Within area **PPc**, extensive and dense reciprocal connectivity was observed throughout the entire cortical area. This extensive reciprocal connectivity was also observed throughout area **PPr**, although if, for example, the injection site in **PPc** was located in the lateral portion of this area, the reciprocal connectivity in the medial region of **PPr** was weaker than that observed in the lateral region (Fig. 5). Within areas **17** and **18**, patches of weak anterograde projections were observed, and while these patches sometimes contained retrogradely labeled cells, the distribution of anterograde and retrograde label was mostly independent of each other.

**Figure 5:**
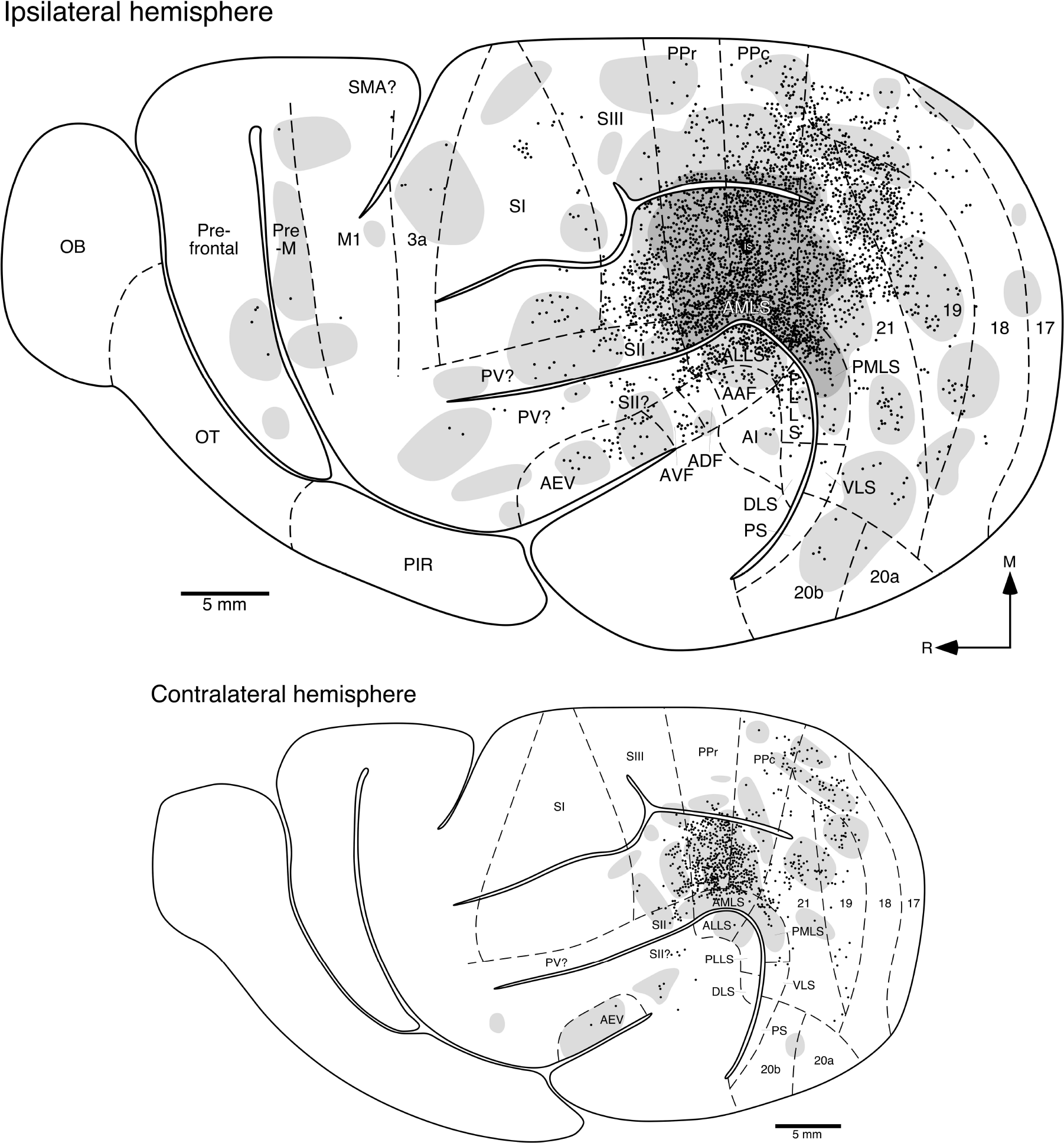
Location of retrogradely labelled cortical neurons (**filled circles**) and anterogradely labelled axons and axon terminals (dense labelling in the **darker grey shading**, light labelling in the **lighter grey shading**) following transport from the injection site (**is**) located in the lateral portion of area **PPc**. The upper larger image represent the distribution of cells and axons in the ipsilateral semi-flattened cerebral hemisphere, while the lower smaller image represents the semi-flattened cerebral hemisphere contralateral to the injection site. Note that ipsilaterally, extensive connectivity is seen through the occipital visual areas (**19** and **21**), the suprasylvian visual areas (**AMLS**, **ALLS** and **PMLS**) as well as the third somatosensory area (**SIII**). In addition, broad, but weaker, reciprocal connectivity is observed in the temporal visual and auditory regions, the rostral somatosensory regions, motor, pre-motor and prefrontal regions, as well as the claustrocortical regions. The contralateral connectivity is much weaker, and less widespread, but still extensive. Areal boundaries were demarcated using alternative sections stained for cytochrome oxidase and the boundaries represent approximations based on this stain and available maps of the ferret brain (Manger, Kiper, Masiello, Murillo, Tettoni, Hunyadi, & Innocenti, 2002a; Manger et al., 2002b; Manger, Nakamura, Valentiniene, & Innocenti, 2004; Manger, Engler, Moll, & Engel, 2005; Manger, Engler, Moll, & Engel, 2008; Manger, Restrepo, & Innocenti, 2010; Bizley, Nodal, Nelken, & King, 2005; Homman-Ludiye et al., 2010). See list for abbreviations.

In contrast, in areas **19** and **21**, the patches of anterograde label were larger and covered the majority of each of these cortical areas, and these patches often contained moderate densities of retrogradely labeled cells. The density of labeling in areas **19** and **21** was always stronger in their more medial aspects, which lie closer to the injection sites, but to reiterate, connectivity was observed through both cortical areas (Fig. 5). Of the suprasylvian visual regions, strong reciprocal connectivity was observed throughout area **AMLS** and the medial half of **PMLS**, with weaker reciprocal connectivity in the lateral half of **PMLS**. Moderately dense reciprocal connectivity was observed throughout area **ALLS**, while weak reciprocal connectivity was observed in area **PLLS**, with areas **DLS** and **VLS** only having weak retrograde connectivity (Fig. 5). Within the visual temporal areas, weak reciprocal connectivity was observed in areas **20a** and **20b**, with area **PS** being devoid of connections.

All somatosensory cortical areas exhibited reciprocal connections with area **PPc**. The densest reciprocal connectivity was observed in area S**III**, with patches of connectivity found throughout the area, but as with area **PPr**, if the injection site in **PPc** was located in the lateral portion of the area, the connectivity in area **SIII** was strongest in the lateral portion and weaker in the medial portion (Fig. 5). Areas **SII**, **PV**, **SI** and **3a** all exhibited patches of weak anterograde connectivity throughout much of their extent, and these patches were often coincident with patches of retrogradely labeled neurons, although often the anterograde and retrograde connections were not matching. In the primary motor (**M1**) and premotor regions, patches of anterograde projections, with occasional retrogradely labeled neurons were observed, with the anterograde projections occupying much of the mediolateral extent of the premotor cortical region (Fig. 5). Similarly, weak patches of anterograde projections, with occasional retrogradely labeled cells were observed in the lateral half of the orbital gyrus, presumably representing the prefrontal cortex. As mentioned, connectivity was observed in the anterior and middle portions of the ectosylvian gyrus, but not the posterior portion of this gyrus. In the middle ectosylvian gyrus, weak reciprocal connectivity was observed in the primary auditory cortex (**A1**), the anterior auditory field (**AAF**) and the anterior dorsal (**ADF**) and anterior ventral (**AVF**) auditory fields. Similarly, weak reciprocal connectivity was observed in the multimodal anterior ectosylvian visual area (**AEV**), with patches of weak anterograde label being observed rostral to this area in the cortex overlying the claustrum. The strongest ipsilateral retrograde connectivity was observed within **PPc** (N% = 25.93%), with a decrease in connectivity strength in area **21** (N% = 16.92%), followed by areas **18** (N% = 9.78%), PPr (N% = 9.55%), **19** (N% = 6.93%), **AMLS** (N% = 6.70%) and **PMLS** (N% = 4.38%), with minor connections in other cortical areas (Fig. 4, Table 1).

Contralateral cortico-cortical connectivity following tracer injection in **PPc** was substantially weaker and less diffuse than that observed in the ipsilateral hemisphere (Fig. 5). In the contralateral **PPc**, a moderately dense cluster of reciprocal connectivity was found homotopic to the injection site, and this cluster spread throughout much of the contralateral **PPc**; that is, if the injection was placed in the lateral portion of **PPc**, label was also observed in the contralateral medial portion of **PPc**, albeit much weaker than in the homotopic lateral region of **PPc** (Fig. 5). Moderately dense reciprocal connectivity was also observed throughout the contralateral **PPr**, but more restricted in its mediolateral extent than seen for **PPc**. Within the contralateral suprasylvian areas, weak reciprocal connectivity was observed in areas **AMLS** and **PMLS**, while weak anterograde connectivity was observed in **ALLS** and **PLLS**, with a few retrogradely labeled cells being observed in the contralateral **VLS** (Fig. 5). Patches of weak reciprocal connectivity were observed in occipital areas **18**, **19** and **21**, but no label was observed in area **17**. A small patch of weak anterograde connectivity was observed at the border of temporal areas **20a** and **20b**. Occasional retrogradely labeled neurons were observed surrounding these patches of reciprocal connectivity in these three occipital areas. Weak patches of reciprocal connectivity were observed in the somatosensory areas **SIII** and **SII**, with the extent of the anterograde projections being larger than those of the retrograde connections (Fig. 5). Weak reciprocal connectivity was also observed in the multimodal temporal area **AEV**, and in the anterior ventral (**AVF**) and anterior dorsal (**ADF**) fields of auditory cortex, with a small patch of weak anterograde label found rostral to these regions in the cortex overlying the claustrum. Thus, the connectivity between the hemispheres is far weaker than that within the hemisphere (Table 1, Figs. 4, 5), and these contralateral connections are substantially less widespread than the ipsilateral connections following injections of tracer into **PPc**.

Following injection of tracer into **PPc**, widespread, mostly reciprocal, connectivity was observed in specific nuclei of the visual and motor portions of the dorsal thalamus (Figs. 3a, 6). Within the lateral geniculate nucleus, no anterograde labeling was observed, but retrogradely labeled neurons were seen sparsely distributed throughout lamina **A/A1**, lamina **C** and in the **MIN**. Thus, while the lateral geniculate nucleus does project to **PPc**, it does not receive projections from this cortical area. Both the lateral posterior and pulvinar visual thalamic nuclei were reciprocally connected with area **PPc** (Fig. 6). In both nuclei large, often dense, patches of anterograde label were observed throughout the rostrocaudal extent of these nuclei. Within these patches of anterograde label a moderate density of retrogradely labeled cells were also observed. In addition, low densities of retrogradely labeled cells were observed beyond these patches of reciprocal connectivity. At more rostral levels, substantial patches of reciprocal connectivity were observed throughout the ventral anterior (**VA**) nucleus, part of the motor system, and as with the connectivity of the lateral posterior and pulvinar nuclei, low densities of retrogradely labeled cells were found surrounding these patches of reciprocal connectivity (Fig. 6a-c). Thus, **PPc** appears to be most strongly reciprocally connected with the lateral posterior, pulvinar (visual) and **VA** (motor) nuclei of the dorsal thalamus.

**Figure 6:**
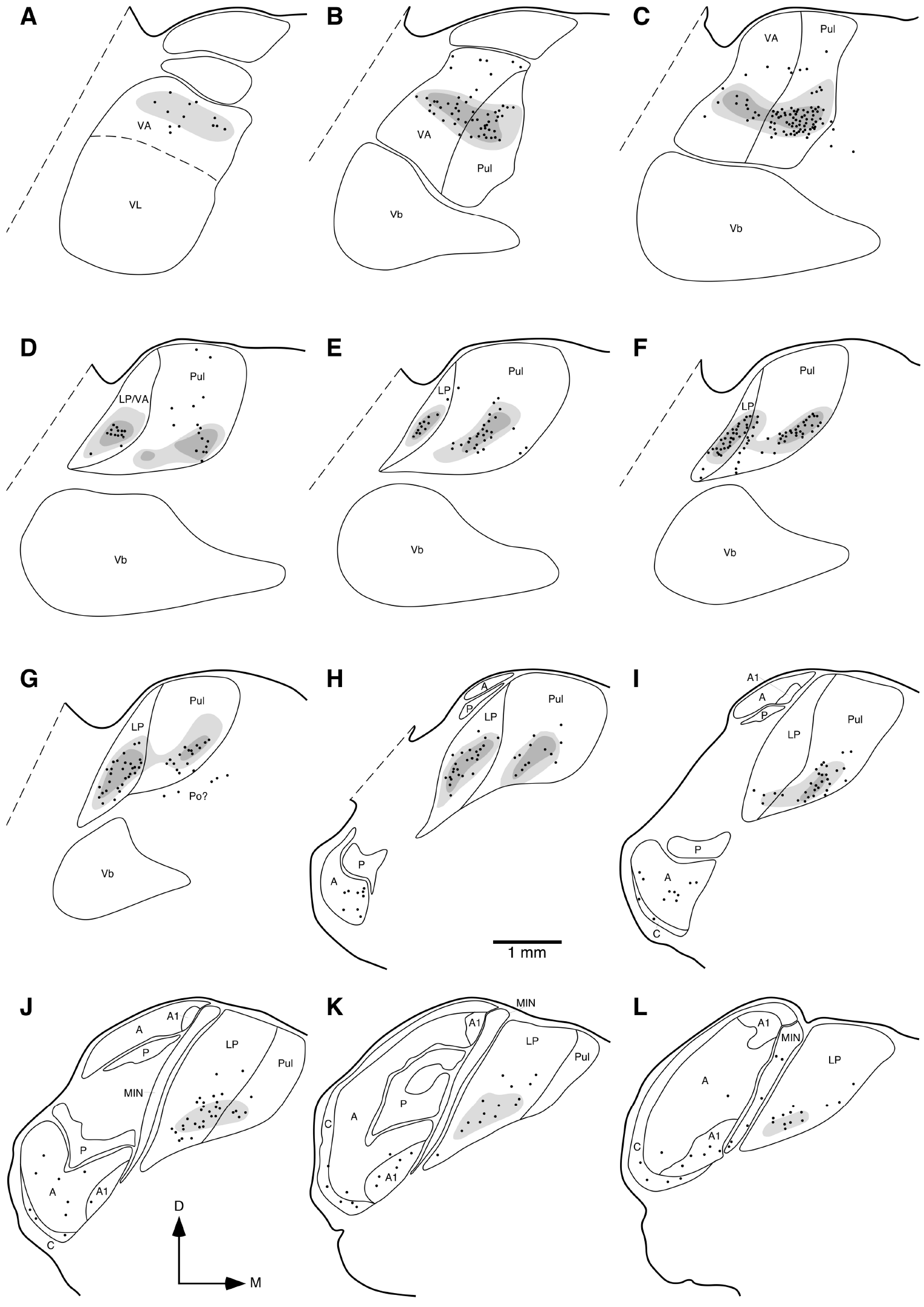
Diagrammatic reconstructions of the location of retrogradely labelled cells (**filled circles**) and anterogradely labelled axons and axon terminals (dense labelling in the **darker grey shading**, light labelling in the **lighter grey shading**) in the visual and motor thalamus of the ferret following injection of tracer into the lateral portion of the parietal visual area **PPc**. **a**represents the most rostral coronal section, with **l**being the most caudal. Each section is approximately 400 μm apart. Note the dense reciprocal connectivity with the lateral posterior (**LP**), pulvinar (**Pul**) and ventral anterior (**VA**) nuclei, and limited label in the lateral geniculate nucleus. In all images dorsal (**D**) is to the top and medial (**M**) to the right. See list for abbreviations.

### Connectivity of PPr

As with area **PPc**, injection of tracer into **PPr** revealed widespread connectivity throughout the vast majority of the ipsilateral cerebral cortex (Fig. 7). The only regions of the cerebral cortex where connectivity was sparse were areas **17**, **18**, and the middle and posterior portions of the ectosylvian gyrus (Fig. 7). Within area **PPr**, extensive and high density reciprocal connectivity was observed throughout this cortical area, but if, for example, the injection was made in the medial portion of **PPr**, the strongest connectivity was observed in the medial portion, with somewhat weaker connectivity observed in the lateral portion (Fig. 7). Strong reciprocal connectivity was observed throughout area **PPc** following tracer injection into area **PPr**, with again the differential between medial and lateral portion connectivity strength based on the location of the injection site. Within area **17** only the occasional weak patch of anterograde projections was observed, while in area **18** slightly more weak patches of anterograde projections, associated with a small number of retrogradely labeled cells, was observed. The extent of the connectivity increased progressively in areas **19** and **21**, with the majority of area **21** showing weak to moderate reciprocal connectivity with area **PPr** (Fig. 7). Within the suprasylvian visual areas, weak to moderate reciprocal connectivity was observed throughout areas **AMLS** and **ALLS**, with areas **PMLS** and **PLLS** showing similar connectivity in the medial half of these areas. A weak patch of anterograde projections was observed in area **DLS**, while area **VLS** was devoid of any connections to **PPr**. Of the temporal visual areas, both areas **20a** and **PS** showed no connections with **PPr**, and a weak patch of anterograde projections was observed in area **20b** and in the ventral posterior ectosylvian region (**VP**), this being the only region of the posterior ectosylvian gyrus showing connectivity to **PPr** (Fig. 7).

**Figure 7:**
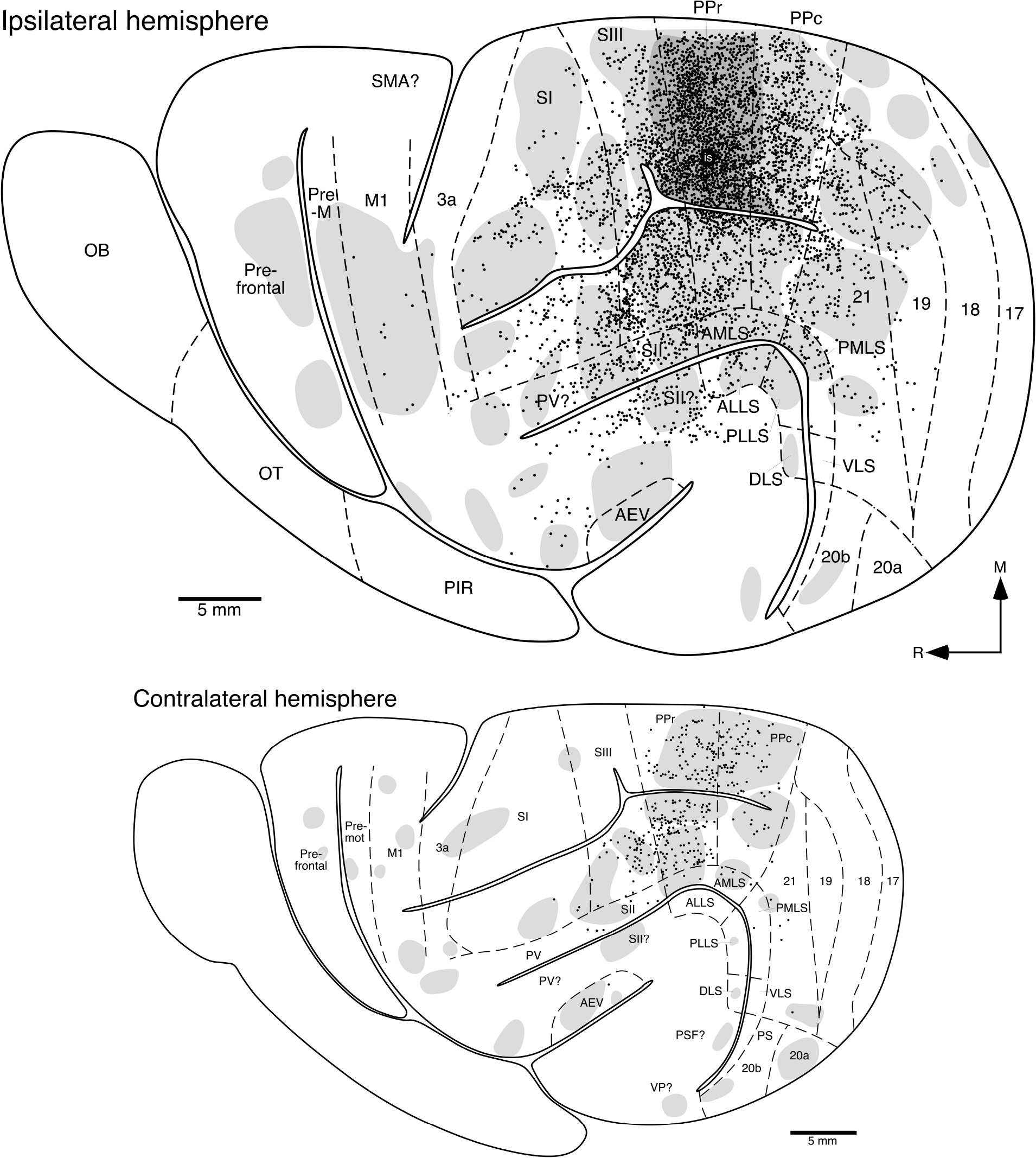
Location of retrogradely labelled cortical neurons (**filled circles**) and anterogradely labelled axons and axon terminals (dense labelling in the **darker grey shading**, light labelling in the **lighter grey shading**) following transport from the injection site (**is**) located in the medial portion of area **PPr**. All conventions and abbreviations as provided in the legend to Fig. 5. Note that ipsilaterally, extensive connectivity is seen through the occipital visual areas (**19** and **21**), the somatosensory areas (**SI**, **SII**, **SIII** and **PV?**). In addition, broad, but weaker, connectivity is observed in the suprasylvian visual area, the motor, pre-motor and prefrontal regions, as well as the claustrocortical regions. The contralateral connectivity is much weaker, and similarly widespread. See list for abbreviations.

All somatosensory cortical areas showed connectivity to area **PPr**, but this was very weak in area **3a**. The most extensively connected somatosensory area was area **SIII**, which showed high to moderate densities of reciprocal connectivity throughout. Moderate to low-density reciprocal connectivity was observed in areas **SI**, **SII/PV** (Fig. 7). A large and broad weak anterograde projection was observed throughout much of the primary motor cortex (**M1**) and the premotor cortical region, although only a few retrogradely labeled neurons were observed in **M1**, and no retrogradely labeled cells were observed in the premotor region. Patches of weak anterograde projections were observed in the caudal half of the orbital gyrus, presumably the prefrontal cortex. Within the middle ectosylvian gyrus, small clusters of retrogradely labeled neurons were observed in the anterior auditory field (**AAF**), while in the anterior ectosylvian gyrus weak anterograde projections were observed in the multimodal area **AEV**. Rostral to the anterior ectosylvian gyrus, in the cortex overlying the claustrum, mismatched patches of weak anterograde and retrograde label was observed (Fig. 7). The strongest ipsilateral retrograde connectivity was observed within area **PPr** (N% = 27.32%), with a decrease in connectivity strength in **PPc** (N% = 19.43%) and SIII (N% = 17.09%), followed by areas **21** (N% = 6.92%), **SI** (N% = 6.20%), **SII/PV** (N% = 4.09%), **AMLS** (N% = 2.62%) and **M1** (N% = 2.59%), with minor connections in other cortical regions (Table 1, Fig. 4).

Contralateral cortico-cortical connectivity following tracer injection in **PPr** was substantially weaker and less diffuse than that observed in the ipsilateral hemisphere (Fig. 7). In the contralateral **PPr**, moderately dense reciprocal connectivity was observed throughout this cortical area. Such reciprocal connectivity was also found throughout area **PPc**, although the numbers of retrogradely labelled neurons was higher in the region corresponding to the injection site, that is, if the injection was made in the medial portion of **PPr**, the number of retrogradely labelled neurons was higher in the medial portion of the contralateral **PPc** (Fig. 7). Weak reciprocal connectivity was observed in the suprasylvian visual areas **AMLS** and **PMLS**, with small patches of weak anterograde projections being observed in **PLLS** and **DLS**. Weak contralateral reciprocal connectivity was only observed in area **21** of the occipital visual areas, with weak patches of anterograde connectivity observed in the temporal visual areas **20a**, **20b** and **AEV**, as well as in the posterior suprasylvian auditory field (**PSF**) and the multimodal ventral posterior ectosylvian region (**VP**). Weak, but broad, reciprocal connectivity was observed with somatosensory area **SIII**, while weak patches of reciprocal connectivity were observed in areas **SII** and **SI**. Patches of weak anterograde label were observed in areas **3a**, primary motor cortex (**M1**), premotor cortex and the orbital gyrus (presumably prefrontal cortex) (Fig. 7). Lastly, isolated patches of weak anterograde projections were observed in the cortex overlying the claustrum. Thus, while the connectivity between the hemispheres is far weaker than that within the hemisphere (Table 1, Figs. 4, 7), these contralateral connections are nearly as widespread as the ipsilateral connections following injections of tracer into **PPr** (Fig. 7).

Following injection of tracer into **PPr**, widespread, mostly reciprocal, connectivity was observed throughout specific nuclei of the visual, motor and somatosensory regions of the dorsal thalamus (Figs. 3b, 8). A small number of retrogradely labeled neurons was observed in the **A/A1** and **C** lamina and **MIN** of the lateral geniculate nucleus. A small patch of reciprocal connectivity was observed in the lateral posterior nucleus, with occasional retrogradely labeled neurons found surrounding this small patch of reciprocal connectivity (Fig. 8j-l). A dense patch of reciprocal connectivity was observed throughout the rostrocaudal extent of the pulvinar nucleus, with occasional retrogradely labeled cells surrounding this patch of reciprocal connectivity. Further rostral within the thalamus, a large and dense patch of reciprocal connectivity was observed throughout the majority of the ventral anterior (**VA**) nucleus of the motor portion of the ferret dorsal thalamus (Fig. 8a-i), but no label was observed in the ventral lateral (**VL**) nucleus of this region of the thalamus. Occasional retrogradely labeled neurons were observed in the ventrobasal complex (**Vb**), the medial division of the posterior nucleus (**Po**), and the anterior ventral nucleus (**AV**) (Fig. 8). Thus, **PPr** appears to be most strongly reciprocally connected with the pulvinar (visual) and ventral anterior (**VA**, motor) nuclei of the dorsal thalamus.

**Figure 8:**
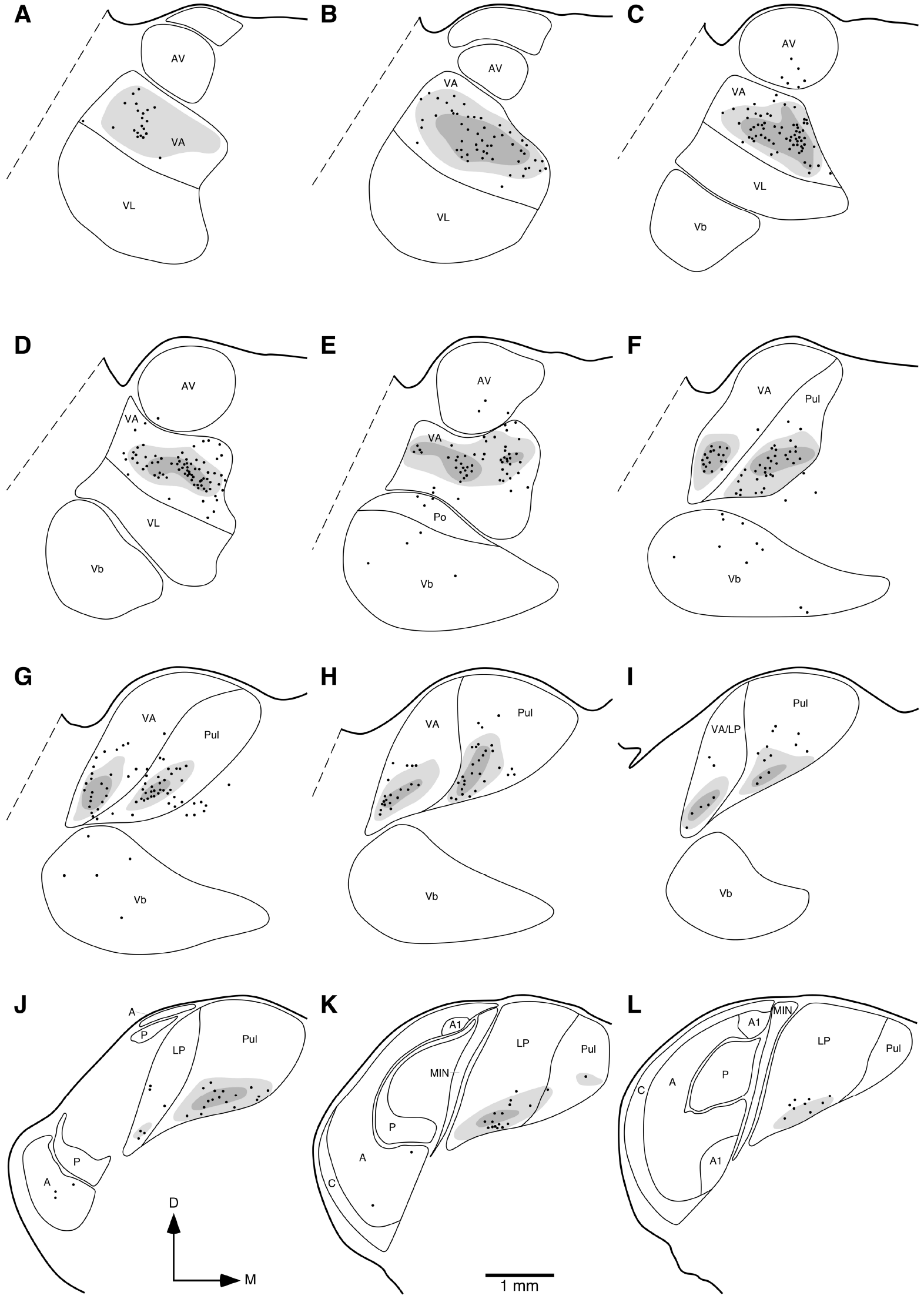
Diagrammatic reconstructions of the location of retrogradely labelled cells (filled circles) and anterogradely labelled axons and axon terminals (dense labelling in the **darker grey shading**, light labelling in the **lighter grey shading**) in the visual and motor thalamus of the ferret following injection of tracer into the medial portion of the parietal visual area **PPr**. **a**represents the most rostral coronal section, with **l** being the most caudal. Each section is approximately 400 μm apart. Note the dense connectivity with the pulvinar nucleus of the visual thalamus and the ventral anterior (**VA**) nucleus of the motor thalamus, with some weak connectivity with the lateral geniculate nucleus and the ventrobasal nuclei. Conventions and abbreviations as provided in the legend to Fig. 6.

## Discussion

The present paper examined the ipsilateral and contralateral cortico-cortical and corticothalamic connectivity of the posterior parietal cortical areas **PPc** and **PPr** in the ferret, by means of standard anatomical tract-tracing methods. The data provides an understanding of the connectivity of the posterior parietal cortex of the ferret, which will contribute to the Ferretome connectivity database (www.ferretome.org; Sukhinin, Engel, Manger, & Hilgetag, 2016), facilitate cross-species analyses of connectomes, and provide insights into proposed principles of cortical connectivity and cortical area evolution. Although **PPc** and **PPr** are strongly interconnected, it was observed that **PPc** was more strongly connected with the occipital and temporal visual cortex than **PPr**, while **PPr** was more strongly connected with the somatosensory cortex than **PPc** (Figs. 4, 5; Table 1), reflecting their specific functional aspects (Manger et al., 2002b). For all **PPc** and **PPr** injection cases, it was evident that the ipsilateral hemisphere was more strongly connected than the contralateral hemisphere, but that the distribution of connections in the contralateral hemisphere mirrored that of the ipsilateral hemisphere (Figs. 4, 5, 7; Table 1) similar to what has been observed in connectional studies of this region in the macaque monkey, rat and mouse (Barbas, Hilgetag, Saha, Dermon, & Suski, 2005; Goulas, Uylings, & Hilgetag, 2017; Swanson, Hahn, & Sporns, 2017). Numerous connectional similarities and differences were observed between the posterior parietal cortex of the ferret and the cat, and the implications of these differences are discussed. Unfortunately, our understanding of the posterior parietal cortical region in the cat is incomplete (Heath & Jones, 1971; Marcotte & Updyke, 1982; Symonds & Rosenquist, 1984; Olson & Lawler, 1987; Pigarev & Rodionova, 1998), but from the available studies it appears reasonable, for comparative purposes, to suggest that ferret areas **PPc** and **PPr** are homologous or analogous to areas **7** and **5** of the cat, respectively; however, we suggest this cautiously and use this comparison as a working hypothesis until confirmation or refutation is obtained.

### Area PPc connectivity − ferret vs cat

Ferret area **PPc** displayed extensive ipsilateral cortical connectivity, being connected with all 15 visual areas previously described for the ferret (Fig. 9), although the strongest connectivity was with **PPr**, occipital visual areas **18**, **19** and **21**, and suprasylvian visual areas **PMLS** and **AMLS**. In addition, **PPc** of the ferret was connected with all known somatosensory and motor areas of the ferret, most strongly with areas **SIII**, **SII/PV** and **SI**. When compared to studies of the connectivity of area **7** of the cat, while the cat shows a similar overall connectional topography (Kawamura, 1973; Babb, Waters, & Asanuma, 1984; Symonds & Rosenquist, 1984; Olson & Lawler, 1987; Avendaño, Rausell, Perez-Aguilar, & Isorna, 1988; Cavada & Reinoso-Suárez, 1985; Scannell, Blakemore, & Young, 1995), the ferret **PPc** appears to have a far broader range of connections to other visual areas than observed in the cat (Fig. 9), the cat lacking connections to areas **17**, **PMLS**, **VLS**, and **AES**; however, all of these connections, apart from the **PPc-PMLS** connection observed in the ferret, are weak connections. Thus, the differences observed may be due to methodology, although the strong **PPc-PMLS** connection present in the ferret, but absent in the cat, is a difference of note. Interestingly, the ipsilateral connectivity to the somatosensory cortex of ferret **PPc** and cat area **7** appear very similar, the one difference being the presence of a weak connection to **SI** in the ferret, which is absent in the cat (Fig. 9; Jones & Powell, 1968; Scannell et al., 1995). The interhemispheric connectivity of ferret **PPc** was more restricted than its ipsilateral connectivity, although still quite broad. Significant callosal connections were observed with contralateral areas **PPc**, **PPr**, **18**, **19**, **21**, **PMLS**, **AMLS**, **SIII** and **SII**, with weaker connections to other cortical areas, but notably absent from motor and premotor areas (Figs. 4, 5). Studies of the interhemispheric connectivity of area **7** in the cat (possible homologue to area **PPc** in ferret) are limited, but the consensus is that the callosal connectivity is more restricted than the ipsilateral connectivity, a feature observed in the ferret as well as all mammalian species studied to date (Swanson et al., 2017). Furthermore, area **7** in the cat is similarly connected to its homotopic area as well as to the heterotopic area **5** (Garol, 1942; Heath & Jones, 1971; Kawamura, 1973) (possible homologue to **PPr** in ferret), indicating that the posterior parietal visual regions in the ferret and the cat show similar patterns of callosal connectivity.

**Figure 9:**
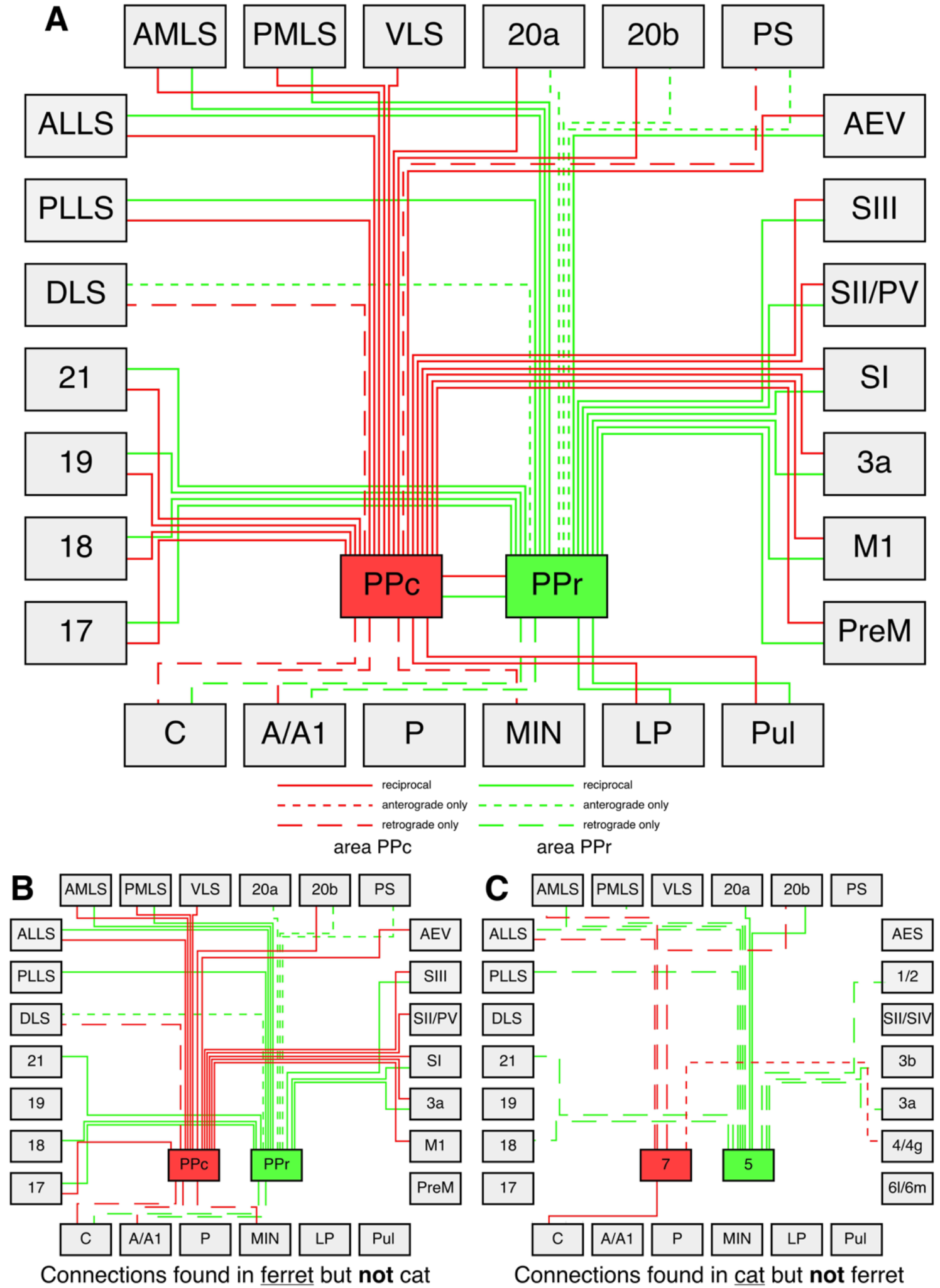
Wiring diagrams depicting the connectivity of areas **PPc** and **PPr** with each other, other visual cortical areas ipsilaterally, and the visual thalamus in the ferret (**a**), the connections of these areas observed in the ferret but not the cat (**b**), and the connections of these areas overs in the cat but not the ferret (**c**). Each colour represents a specific cortical area (**PPc**- red, **PPr**- green), with solid lines representing reciprocal connections, short dashed lines anterograde connections only, and long dashed lines retrograde connections only. (**a**) Of the potential 156 connections (for each cortical area 26 potential reciprocal, anterograde only or retrograde only connections) of these two areas to the different visual cortical and thalamic regions, a total of 50 connections are found in the ferret. (**b**) In the ferret 32 connections not found in the cat were observed, while (**c**) in the cat 16 connections not found in the ferret were observed (Garol, 1942; Jones & Powell, 1968; Heath & Jones, 1971; Kawamura, 1973; Robertson & Cunningham, 1981; Raczkowski & Rosenquist, 1983; Babb et al., 1984; Symonds & Rosenquist, 1984; Avendano et al., 1988; Scannell et al., 1995, 1999).

In terms of thalamic connectivity, area **PPc** of the ferret was reciprocally connected with the **LP** and pulvinar nuclei and received projections from the **A**, **C** and **MIN** laminae of the **LGN**. While the connections of cat area **7** with the **LP** and pulvinar are similar to that observed in ferret **PPc**, cat area **7** was reciprocally connected with lamina **C** of the **LGN**, but lacked connections with laminae **A** and **MIN** (Fig. 9; Heath & Jones, 1971; Graybiel, 1972; Hendry, Jones, & Graham, 1979; Robertson & Cunningham, 1981; Raczkowski & Rosenquist, 1983; Olson & Lawler, 1987; Scannell, Burns, Hilgetag, O’Neil, & Young, 1999). Thus, when comparing the global connectivity of ferret area **PPc** with cat area **7**, it appears that the connectivity patterns of the ferret and cat are generally similar, but that the ferret has a more broadly connected posterior parietal unimodal visual area than the cat. While the observed differences may be methodological (e.g. different tract tracers used), there are certain differences that are unlikely to be explained methodologically, specifically the connectivity of **PPc** with **PMLS** and the various laminae of the **LGN** in the ferret. In our previous study of the connectivity of the occipital visual area of the ferret (Dell, Innocenti, Hilgetag, & Manger, 2018), we noted that **PMLS** appears to form a hub in the network processing visual information in the ferret, and this is supported in the current study. In addition, we noted that when comparing the connectivity of the visual cortex with the **LGN** in the ferret and cat, that the **LGN** appears to be more strongly connected with visual areas early in the processing hierarchy in the cat, whereas the **LGN** is more strongly connected with visual areas later in the hierarchy in the ferret, and this pattern appears to continue into the posterior parietal cortex.

### Area PPr connectivity − ferret vs cat

Ferret **PPr**, like **PPc**, was broadly connected with almost all ipsilateral visual cortical areas, **VLS** being the only area not connected with **PPr** (Fig. 9), but the strongest connections to other ipsilateral visual areas were with **PPc**, **21** and **AMLS**. In addition, **PPr** was connected with all known ipsilateral somatomotor areas of the ferret, with strongest connectivity observed with areas **SIII**, **SI**, **SII/PV** and **M1**. When compared with area **5** of the cat, the majority of these connections show strong similarity (Fig. 9; Jones & Powell, 1968; Kawamura, 1973; Babb et al., 1984; Symonds & Rosenquist, 1984; Aveñdano et al., 1988; Cavada & Reinoso-Suárez, 1985; Scannell et al., 1995), but there are several differences in terms of whether the connections are reciprocal, anterograde only, or retrograde only (Fig. 9). Major differences in the connectivity patterns to visual cortical areas between ferret **PPr** and cat area **5** include, reciprocal connections with area **17** and anterograde projections to area **PS** in the ferret that are absent in the cat (Fig. 9; Kawamura, 1973; Aveñdano et al., 1988; Scannell et al., 1995), and mostly reciprocal connections with the suprasylvian visual areas in the ferret, whereas these are mostly retrograde connections in the cat (Fig. 9; Kawamura, 1973; Aveñdano et al., 1988; Scannell et al., 1995). For connectivity with the somatomotor cortex, the areas connected in the ferret and cat appear broadly similar (Fig. 9; Jones & Powell, 1968; Kawamura, 1973; Avendaño et al., 1988; Scannell et al., 1995), but the ferret shows reciprocal connectivity with areas SI**II**, **SI** and **3a**, whereas these regions only send connections to area **5** in the cat (Fig. 9; Jones & Powell, 1968; Kawamura, 1973; Avendaño et al., 1988; Scannell et al., 1995). The interhemispheric connectivity of ferret **PPr** was more restricted than its ipsilateral connectivity, although still quite broad, especially the anterograde projections. Significant callosal connections were observed with the contralateral areas **PPr**, **PPc** and **SIII**, with weaker anterograde projections to many other cortical areas, including motor, premotor and prefrontal areas (Figs. 4, 7). As observed in the ferret, the interhemispheric connectivity of cat area **5** (potential homologue to ferret area **PPr**) was more restricted than that of the ipsilateral hemisphere and, unlike the ferret, callosal connections were observed with only areas **5** and **7** (Garol, 1942; Heath & Jones, 1971; Kawamura, 1973), indicating that while the posterior parietal areas maintain strong interhemispheric connections in both species, these connections are broader in the ferret than the cat.

In terms of thalamic connectivity, area **PPr** of the ferret was reciprocally connected with the **LP** and pulvinar nuclei and received projections from the **A** and **C** laminae of the **LGN**. While the connections of cat area **5** with the **LP** and pulvinar are similar to that observed in ferret **PPr**, cat area **5** lacked connections with the **LGN** (Fig. 9; Heath & Jones, 1971; Graybiel, 1972; Hendry et al., 1979; Robertson & Cunningham, 1981; Raczkowski & Rosenquist, 1983; Scannell et al., 1999). Thus, as with area **PPc**, area **PPr** of the ferret appears to be more broadly connected with other cortical areas than area **5** of the cat. In addition, **PPr** of the ferret receives projections from the **LGN**, which are not observed in the cat, furthering the pattern observed, where the **LGN** of the ferret is more strongly connected with higher order visual areas in the ferret, but with lower order visual areas in the cat. Despite these differences, the overall connectivity pattern is reasonably consistent between species and certain differences observed may be based in different methodologies. These findings support the notion that a common connectional blue print may exist across species within an order and that differences may be due to evolutionary or functional constraints or specializations.

### Connectivity patterns of parietal areas of the ferret and the cat

There is clearly a global similarity in the patterns of connectivity of posterior parietal areas **PPc** and **PPr** of the ferret and area **5** and **7** of the cat, with ipsilateral and contralateral cortical areas and also with the visual thalamus. Despite this similarity, there are differences of interest that emerge from this comparison. The ferret posterior parietal areas tend to have more reciprocal connections with other cortical areas than the cat, all of which appear to tend towards retrograde connectivity (Fig. 9), indicating that the parietal areas in the cat receive more information than they send. Thus, the ferret posterior parietal cortex may play a greater role in systems level processing and information integration than that of the cat. The second difference is the extended connectivity of the **LGN** to the posterior parietal cortex of the ferret, which appears to be absent in the cat (Fig. 9; Robertson & Cunningham, 1981; Raczkowski & Rosenquist, 1983; Scannell et al., 1999). This underscores a potential emphasis in the ferret on higher order visual stimuli detection compared to lower order visual stimulus detection emphasis in the cat, similar to that seen for the occipital visual areas (Dell et al., 2018). As discussed previously (Dell et al., 2018), these differences may relate to the larger brain size of the cat compared to the ferret, the extensive phylogenetic independence of the two species, anatomical differences related to eye position (lateral in the ferret, frontal in the cat), or their specific strategies for acquiring nutrition. Further studies of species within the two major lineages of the carnivores are required to determine which of these possibilities explains the observed differences.

### Evolution of posterior parietal cortex and the fronto-parietal network

Due to the involvement of posterior parietal cortex in visually-guided motor tasks (Creem-Regehr, 2009), and the small size of this cortical region in most non-primate mammals (Kolb & Walkey, 1987; Wallace et al., 2004; Remple et al., 2006, 2007; Goldring & Krubitzer, 2017; Kaas et al., 2017), studies of the posterior parietal cortex have been mostly undertaken in primates. One of the central themes to emerge from these primate studies is that the posterior parietal cortex is heavily interconnected with motor and premotor cortical areas, forming a distinct reciprocal connectivity network (the fronto-parietal network) that is thought to underlie the visually-guided digital manipulative competencies of primates (e.g. Felleman & Van Essen, 1991; Bullmore & Sporns, 2005; Cloutman & Lambon Ralph, 2012; Caminiti et al., 2015; Kaas & Stepniewska, 2016; Kaas et al., 2017), especially in humans where this region is greatly expanded (Hill et al., 2010). However, the primate studies have led to an extensive parcellation of the posterior parietal cortex (Seltzer & Pandya, 1980; Lewis & Van Essen, 2000a,b; Nelson et al., 2010; Mars et al., 2011; Stepniewska et al., 2011; Kaas et al., 2017), making comparisons to non-primate mammalian species, where this cortical region is not greatly expanded (Goldring & Krubitzer, 2017), difficult. In order to meaningfully compare the organization of primate posterior parietal cortex with that found in other species, the primate with the simplest organizational scheme, the galago (Fang et al., 2005; Stepniewska et al., 2009a,b, 2011; Kaas & Stepniewska, 2016; Kaas et al., 2017), is the best starting point. Organizational schemes forwarded for other larger brained primate species, due to their complexity, are not considered here.

The posterior parietal cortex of the galago has been subdivided into three distinct cortical areas, rostral (visuomotor, heavily connected with motor and pre-motor cortex, **PPCr**), caudal (visual, **PPCc**) and medial (somatovisual, PPCm) (Kaas et al., 2017). While the **PPCm** identified in the galago has no clear counterpart in non-primate mammals, as this midline region has not been explored, **PPCr** and **PPCc** of the galago may be compared with **PPr** and **PPc** of the ferret (Manger et al., 2002b) and PP and OP of the megachiropterans (Rosa, 1999). Due to the lack of distinct posterior parietal cortical areas in most mammals, related to the small size of this region (Goldring & Krubitzer, 2017; Kaas et al., 2017), and the phylogenetic positions of the primates, carnivores and megachiropterans (Arnason, Adegoke, Gullberg, Harley, Janke, & Kullberg, 2008; Foley, Springer, & Teeling, 2016), it would be imprudent to assume that these cortical areas in the galago (**PPCr**, **PPCc**), ferret (**PPr**, **PPc**) and megachiropteran (**PP**, **OP**) are homologous. This indicates that expansion and arealization of the posterior parietal cortex has evolved independently at least three times in mammals; unless one takes the position that the megachiropterans are a sister group to primates (Pettigrew, 1986; Pettigrew, Jamieson, Robson, Hall, McNally, & Cooper, 1989; Pettigrew, Maseko, & Manger, 2008), then it is reasonable to propose homology for areas **PPCc/OP** and **PPCr/PP** in galago and megachiropterans, but not with the ferret, indicating only two currently known independent expansions of the parietal cortex. Despite the intricacies of chiropteran phylogenetic relationships, it is at least reasonable to assume that, at a minimum, expansion and arealization of the posterior parietal cortex has occurred independently in certain lineages within the Euarchontoglires (primates and/or megachiropterans) and Laurasiatheria (carnivores and/or megachiropterans). That there are broad organizational similarities, in that there is a rostral somatovisual area and caudal visual area in all species that have an expanded posterior parietal cortical region regardless of their phylogenetic affinities, indicates that the arealization of this region is likely related to underlying functional and connectional aspects of this cortical territory.

Indeed, even in rodents and tree shrews, where the expansion and arealization of posterior parietal cortex has not occurred, indicated by the presence of a single multimodal posterior parietal region immediately caudal to somatosensory cortex (Goldring & Krubitzer, 2017; Kaas et al., 2017), this multimodal region is connected with the motor cortex, forming the frontoparietal network. Thus, it is reasonable to conclude that the fronto-parietal network is a shared feature of all species within the Euarchontoglires and Laurasiatheria, but varies in its degree of expression. Across species, the posterior parietal cortex forms an integral component of the dorsal visual processing stream (Goodale & Milner, 1992; Goodale, 1998; Kaas et al., 2011). Although this processing stream was initially identified as the #x2018;where’ stream, as it deciphers spatial information about objects (Goodale & Milner, 1992; Goodale, 1998), studies in primates proposed that this stream should rather be called the &#x2018;how&#x2019; stream, as its functions extend to coordinating and controlling multisensory motor operations within a spatial attentional context (Goodale & Milner, 1992; Burish, Stepniewska, & Kaas, 2008; Stepniewska et al., 2009b; Kaas et al., 2011). The projections of this ‘how’ stream are congruent with the fronto-parietal network and thus an extension of the dorsal visual processing stream (see Goodale & Milner, 1992; Goodale, 1998). Our studies also identify this fronto-parietal network in the ferret in line with connectivity observed in other non-primate species such as the cat (Heath & Jones, 1971; Cavada & Reinoso-Suárez, 1985), rat (Beckstead, 1979), guinea pig (Pritzel & Markowitsch, 1981) and tree shrew (Remple et al., 2006, 2007), although this fronto-parietal network appears to be more limited in these non-primate species. These comparative observations indicate that the ferret is a potential experimental model to understand the evolution and function of the posterior parietal cortex and the fronto-parietal network across mammals. Further studies, using long-train intracortical microstimulation in the ferret, determining whether the ferret posterior parietal cortex shows a similar mosaic organization of movement domains embedded within the gross topographic organization of this region (Manger et al., 2002b), will further our understanding of this region, not only in the ferret, but in phylogenetically and functionally relevant senses.

## Acknowledgements

The authors wish to thank Mrs. Sonata Valentiniene for her consistently high-quality histological preparations.

## Role of Authors

PRM and GI designed the study and undertook the experimental aspects of the study. LAD, CCH and PRM analyzed the material and LAD wrote the first draft of the paper, which was subsequently edited by GMI, CCH and PRM. All authors had full access to all of the data in the study and take responsibility for the integrity of the data and the accuracy of the data analysis.

